# Intra-Complex Differential Transcription Strategies for Scaffold versus Effector Proteins

**DOI:** 10.64898/2026.06.12.731977

**Authors:** Shuangmei Tian, Ziyu Zhao, Meharie G. Kassie, Emmanuel Annan, Fangyuan Zhang, Chenghang Zong, Beibei Ren, Degeng Wang

## Abstract

Scaffold-organized multi-protein complexes are major drivers of cellular regulatory pathways, but how the expression of their constituent genes is coordinated to support efficient assembly and cell-to-cell homeostasis remains poorly understood. Using snapTotal-seq data, this study addresses this issue at the transcription level through comparative analyses of stochastic transcription bursting – a major driver of expression level and primary source of its intrinsic cell-to-cell noise. We compared scaffold proteins and their effector partners in major regulatory multi-protein complexes: TNRC6A/B/C (in the miRISC complex) and CNOT1/2 (in the CCR4-NOT complex) in mRNA regulation; and AXIN1, APC, KSR2, AKAPs, DLG1, PATJ, HOMER1/2, and SQSTM1 in signal transduction. Our analyses reveal a shared strategy: compared to genes for active effector partner proteins and the general transcriptome, genes for structural scaffold proteins consistently utilize more frequent and smaller-sized bursting to achieve the fine tuning of a homeostatic expression level. This finding establishes a fundamental link between a protein’s physical role within a regulatory complex and the transcriptional bursting kinetics of its gene.

## Introduction

Multi-protein complexes drive almost all major cellular processes. They non-covalently organize proteins performing interrelated functions together, ensuring that these activities occur in close spatiotemporal proximity, couple efficiently, and proceed as a single functional module. Arguably the best-known examples are the electron transfer chain (complexes I to IV) and the ATP synthase complex (complex V) within the oxidative phosphorylation system. Consequently, the expression of the constituent genes must be precisely coordinated to ensure assembly efficiency and cell-to-cell homeostasis.

There are key architectural differences between regulatory and metabolic multi-protein complexes. Metabolic complexes, like the oxidative phosphorylation system, are formed largely by self-assembly, and stoichiometric balance is achieved through shared regulatory mechanisms among the subunit genes^1^; in bacteria, they often form polycistronic operons^2,3^. Regulatory complexes, on the other hand, adopt a different architectural strategy^4^. Their assembly often utilizes dedicated, non-enzymatic scaffold proteins—such as AXIN1 in WNT signaling and TNRC6A/B/C in miRNA-mediated regulatory activity—to spatiotemporally organize their effector partners, such as catalytic enzymes and transcription factors^4^. These scaffolds act as rate-limiting factors, and their precise availability is critical for signaling and regulatory fidelity. However, whether and how cognate gene expression coordination strategies have evolved to support the assembly and function of these scaffold-organized complexes remain a fundamental question in systems biology.

We previously observed differential control of cell-to-cell expression noise for TNRC6A/B/C (the scaffold) and AGO1/2/3 (the effector) within the miRNA-induced silencing complex (miRISC)^5^. Briefly, to study the miRISC self-feedback loop^6,7^, we quantified cell-to-cell gene expression noise at both nascent and mature RNA levels. While TNRC6A/B/C exhibits reduced noise early at the nascent RNA level, AGO1/2/3 achieves noise reduction primarily via post-transcriptional mechanisms^5,8^. In this study, we demonstrate that TNRC6A/B/C and AGO1/2/3 utilize distinct stochastic transcription bursting strategies and show that this pattern is shared across other major scaffold-organized regulatory complexes.

## Materials and Methods

### SnapTotal-seq dataset and analysis

The HEK293T cell snapTotal-seq dataset is publicly available at the NCBI GEO database (accession number GSE202126). Data processing followed the procedure described in the original publication^9^. Briefly, next generation sequencing (NGS) reads were mapped to the human genome assembly (GRCh37) using the STAR aligner (version 2.5.3a). Uniquely mapped reads were assigned to the GENCODE gene annotation (version 19) with the htseq-count software. Nascent and mature RNA reads were distinguished by the presence of intronic sequences.

We performed cell-cycle scoring with the mature RNA expression data. Briefly, mature RNA expression data were normalized to total mature RNA read counts per cell (counts per 100,000 mapped reads). We then utilized the Seurat single-cell transcriptome R toolkit^10^ for cell-cycle scoring, which was used in regression to mitigate the effects of cell-cycle heterogeneity in downstream analyses. For details, please see the original publication^9^.

### Quantification of nascent RNA expression level and cell-to-cell noise via Negative Binomial (NB) distribution regression analysis

Stochastic transcriptional bursting results in non-Poisson distributions of nascent RNA copy numbers across cell populations^11-13^. To quantify this cell-to-cell expression noise, various parameters have been employed; our previous study utilized the coefficient of variation (CV)^5^, one of the widely used metrics in the field^14-16^. Supported by experimental results and the two-state telegraph model^17-21^, NB distribution has emerged as the leading gene expression model^21-23^, is used in standard RNA- and scRNA-seq analysis tools^10,24^, and is therefore utilized in this study.

We performed NB regressions on nascent RNA read counts to quantify mean expression levels (*µ*) and cell-to-cell noise/dispersion (θ), using sequence depths as a regression offset and cell-cycle scores as a covariate to mitigate the effect of cell cycle heterogeneity. Subsequently, the dispersion parameter (θ) of the NB model was used as a metric for cell-to-cell expression noise (1/*θ* ∝ *noise*). To provide a comprehensive view of cell-to-cell variability, the NB CV was also calculated (*CV*^2^=1/*µ* +1/*θ*).

Since snapTotal-seq covers full-length RNA, transcriptional unit (TU) lengths contribute to nascent RNA read counts. As previously described^8^, we performed a log_2_(*µ*) versus log_2_(TU length) LOESS regression, so that the mean expression levels of genes with different TU lengths are comparable. The LOESS residuals are used as adjusted mean expression level in downstream analyses.

### Statistical analysis

R (version 4.6.0) was used for all data analysis and visualization. Standard base R functions were used for sample normalization, parameter calculation, and LOESS regression. Robust linear regression (rlm) and NB regression were performed using the rlm and glm.nb functions, respectively, in the MASS package. Analysis of Covariance (ANCOVA) was performed using the Anova function (Type III Sum of Squares (SS)) from the car package applied to a rlm model.

## Results

### Differential nascent RNA cell-to-cell expression noise control for TNRC6A/B/C and AGO1/2/3

We previously identified a divergence in nascent RNA expression noise between the miRISC scaffold (TNRC6A/B/C) and its active effector (AGO1/2/3)^5^. The study modeled nascent RNA expression levels as log-normal (LN) distributions, utilizing the LN CV as the cell-to-cell expression noise parameter. We found that while both components exhibit reduced noise at the mature mRNA level, only the TNRC6 scaffolds show noise suppression early at the nascent RNA level^5,8^. This differential nascent RNA noise control is confirmed by the CV of the NB model in the present study (Fig. 1). As expected^19,25^, a CV-mean dependence (a negative correlation) is observed across the transcriptome; however, within the context of this dependence, TNRC6A/B/C consistently exhibit lower CV values relative to AGO1/2/3 (Fig. 1). These observations prompted us to investigate whether this scaffold-versus-effector noise disparity is a common feature across other major regulatory multi-protein complexes.

**Figure 1.**
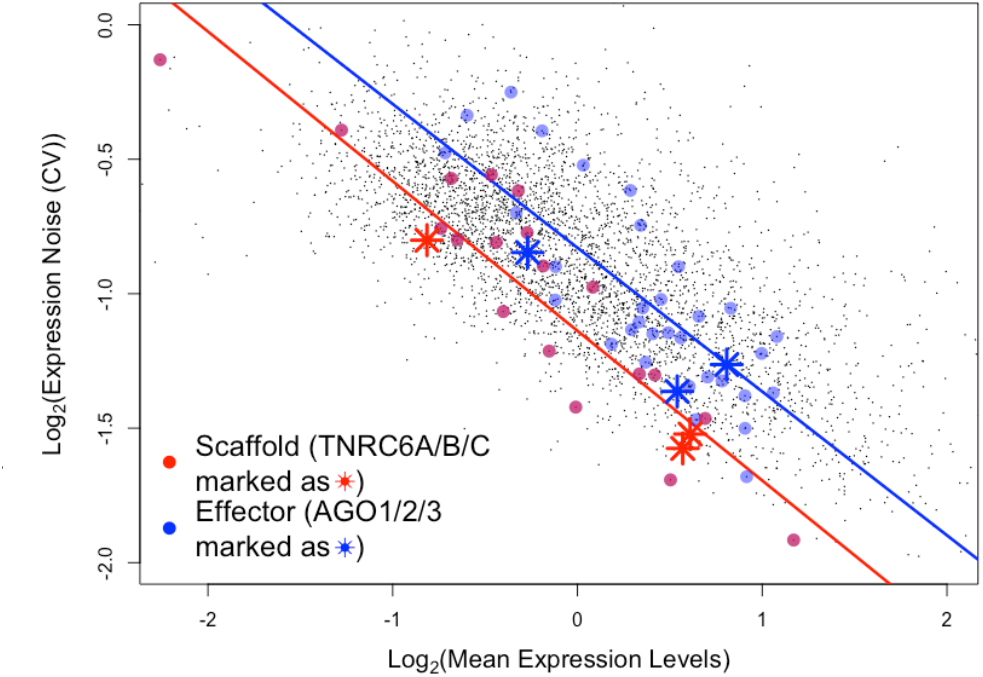
Comparative analysis of cell-to-cell nascent RNA expression noise for scaffold versus effector proteins. Scatterplot of nascent RNA expression noise (measured as the Negative Binomial coefficient of variation, CV) vs. length-adjusted mean expression level. Scaffolds and effectors are highlighted respectively as red and blue datapoints. Linear regression lines illustrate the noise–expression dependence for each group, revealing a significant downward shift in noise for scaffolds across the expression range. Specifically, miRISC complex components TNRC6A/B/C (scaffolds) and AGO1/2/3 (effectors) are marked to highlight the noise divergence.

### Conserved differential scaffold-versus-effector nascent RNA noise control across major regulatory multiple protein complexes

We expanded our analysis to include scaffold proteins with identified effector partners across fundamental regulatory processes (Table 1): AXIN1 and APC in the WNT signaling pathway^26^; KSR2 in the MAP kinase cascades^27^; CNOT1/2 in the CCR4-NOT complex^28^; HOMER1/2 in calcium signaling^29^; DLG1 in T-cell receptor signaling^30^; AKAP1/8/9/10/11/13 as scaffolds for their cognate protein kinase signaling pathways^31^; PATJ and TJP1 in G-protein signaling^32^ and SQSTM1 in autophagy and signal integration^33^. We further included scaffolds essential for vesicle trafficking (AP1G1 and AP1B1)^34^; and the signaling scaffolds HOMER1/2 and DLG1 are also essential for synaptic organization. Highlighted respectively as red and blue datapoints, scaffolds and their partners are well-separated (Fig. 1). As shown by the distinct robust regression (rlm) lines, structural scaffolds consistently exhibit lower cell-to-cell nascent RNA expression noise than their active effector partners (Fig. 1). An ANCOVA confirmed this offset (p-value of 6.05e-5). This suggests that the functional role of a protein as a structural framework—whether providing a platform for signaling or a lattice for cellular architecture—is a primary determinant of its transcriptional burst kinetics.

**Table 1.** List of studied scaffold proteins and their active effector partners detected in the dataset. The major cellular processes or pathways they are involved in are also listed.

| Scaffold | Effector | Process |
| --- | --- | --- |
| TNRC6A/B/C | AGO1/2/3 | MiRNA-mediated regulation |
| CNOT1/2 | CNOT4/6/10 | mRNA de-adenylation, ribosome quality control |
| AXIN1, APC | GSK3B, CSNK1A1, CTNNB1, DYRK1A | WNT signaling |
| KSR2 | RAF1, BRAF, MAP2K1/2, MAPK1/14 | MAP kinase signaling |
| AKAP1, 8-11, 13 | RHOA, CSNK2A1, PRKAR1B, PRKAR2A, PRKCA, PRKD1, PRKD3 | Protein kinase signaling |
| HOMER1/2 | NFATC3, NFAT5, GRID1 | Calcium signaling |
| PATJ, TJP1 | AMOT, DIAPH1, JAM3 | G-protein signaling |
| SQSTM1 | CASP2, PRKCZ, PAWR, RIPK1, GATA4 | Autophagy, signal integration |
| DLG1 | CASK, PRKCA, NET1, AP1S2/3 | Synaptogenesis; T-cell receptor signaling |
| AP1G1, AP1B1 | AP1S2/3 | Subcellular vesicle trafficking |

Next, we explored kinetics of stochastic transcription burst – a major driver of mean expression level and primary source of cell-to-cell noise – for mechanistic into the scaffold-versus-effector protein pattern.

### Differential transcription burst kinetics of scaffold-versus-effector proteins

The Negative Binomial (NB) regression parameters, combined with the foundational two-state telegraph model, enabled a comparative analysis of stochastic transcription burst frequency (ω) and burst size (*s*)^21-23,35^. Human transcription has been experimentally demonstrated to operate within the “bursty limit” (*K*_*off*_ ≫ *K*_*on*_), a regime where the NB model can be utilized to derive burst kinetic parameters and their relationships: ω=*θ* and *s*=*µ*/*θ*. Consequently, the mean expression and burst parameters are linked: *log*_2_(*µ*)=*log*_2_(*θ*) +*log*_2_(*s*).

We performed comparative analyses using these relationships, visualized via a scatter plot of *log*_2_(*µ*) *versus log*_2_(*θ*) (Fig. 2A). The LOESS regression curve represents the transcriptome-wide expected relationship between gene-length-adjusted mean expression and burst frequency; the residuals from this curve quantify *log*_2_(*s*) relative to the global trend. Specifically, positive residuals (points above the regression curve) denote higher burst sizes relative to *log*_2_(*µ*) and *log*_2_(*θ*), while negative residuals denote the opposite. Same as in Figure 1, scaffold and effector proteins are highlighted by red and blue datapoints, respectively (Fig. 2A). Our results clearly demonstrate that scaffold protein genes exhibit lower-than-expected burst sizes, whereas active effector protein genes consistently show the opposite (Fig. 2A). In other words, scaffold protein genes achieve their expression levels via frequent bursts, and active effector protein genes via large bursts.

**Figure 2.**
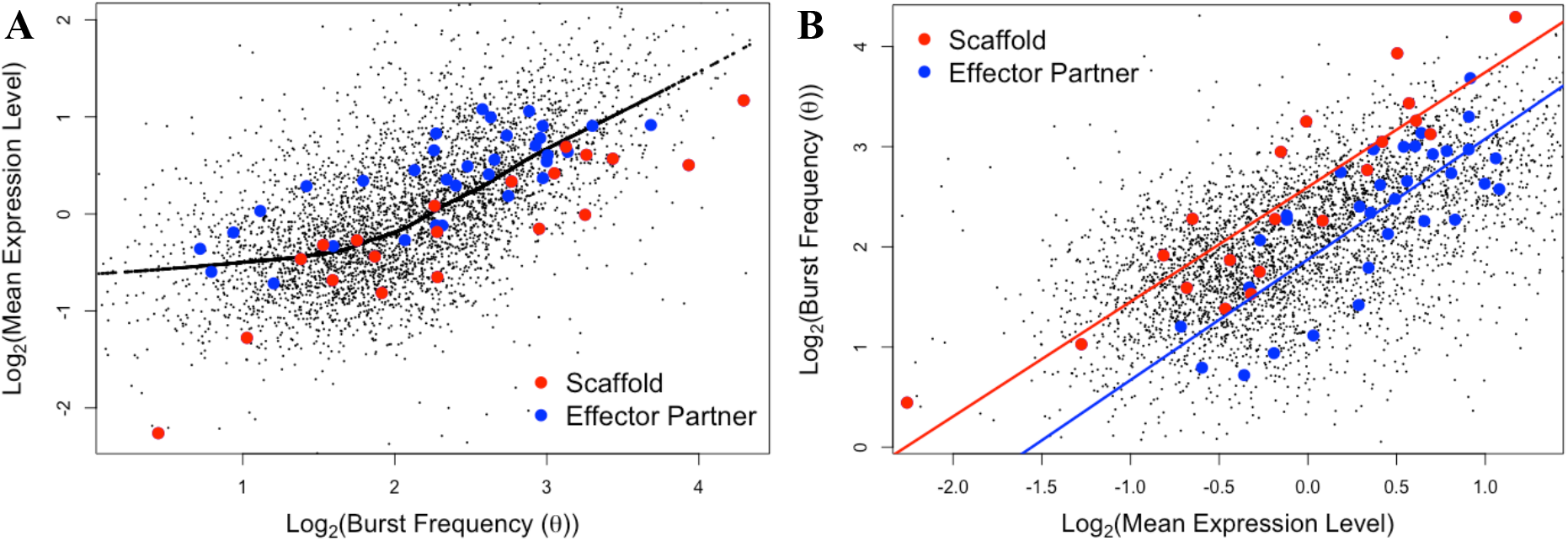
Comparative analysis of transcription burst kinetics for scaffold versus effector proteins. Scaffold and effector protein genes are highlighted as red and blue datapoints, respectively. (**A)** Scatterplot of expression level vs. burst frequency. A LOESS regression curve illustrates the global trend. The residuals relative to this curve serve as a proxy for burst size, showing scaffolds utilize smaller-than-expected bursts compared to effectors. (**B)** Scatterplot of burst frequency vs. expression level. The distinct regression lines for each group illustrate that scaffolds maintain higher burst frequencies than effectors at comparable expression levels.

This divergence is further illustrated in a scatter plot of *log*_2_(*θ*) *versus log*_2_(*µ*) (Fig. 2B). Distinct regression lines for each group illustrate that scaffolds maintain higher burst frequencies than effectors at comparable expression levels (Fig. 2B). An ANCOVA confirmed statistical significance of the divergence (p-value: 4.86e-5).

Finally, the differential transcription strategies are shown directly in a burst size (*s*) versus burst frequency (*θ*) scatter plot (Fig. 3). Burst size and frequency are negatively correlated, as shown by the LOESS regression curve. A clear bifurcated strategy is revealed: scaffolds favoring frequent but small bursts, while effectors utilizing infrequent but large bursts.

**Figure 3.**
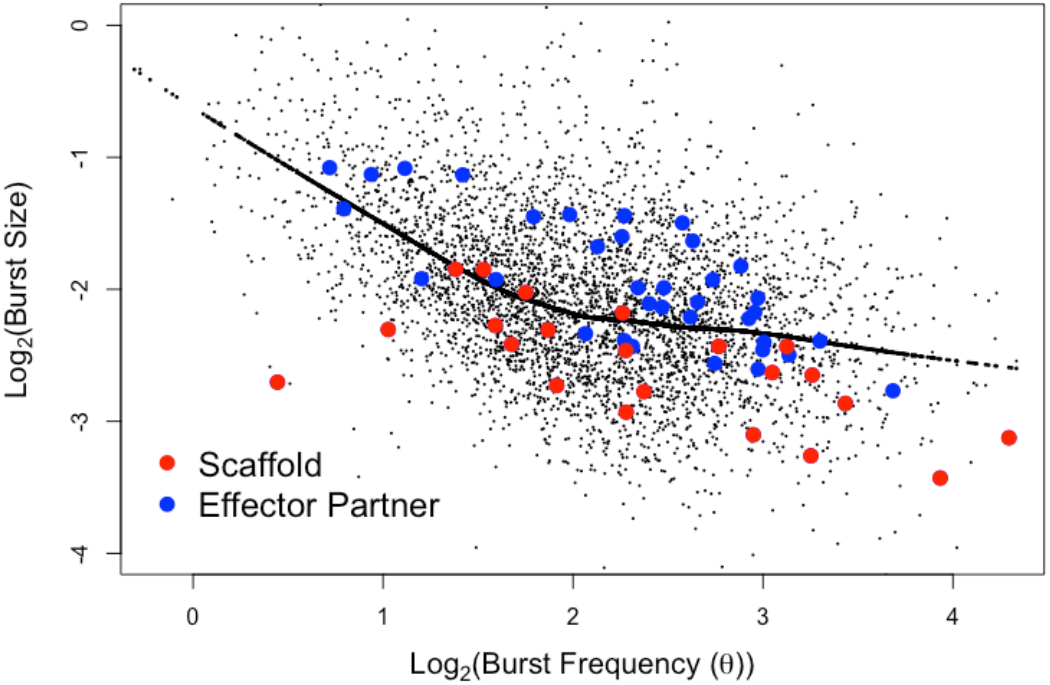
Global landscape of transcription burst size versus frequency. Scatterplot of derived burst size vs. frequency is shown. Scaffold and effector proteins are highlighted as red and blue datapoints, respectively. A LOESS regression curve illustrates the global relationship between the two kinetic parameters. The plot reveals a clear bifurcated strategy: scaffolds favoring a high-frequency, small-burst-size regime, while effectors utilizing an infrequent, large-burst-size regime.

## Discussion

Human gene transcription occurs in stochastic bursts characterized by specific frequencies and sizes. These bursts exhibit function-specific profiles; for example, ribosomal protein genes utilize frequent but relatively small-sized bursts to maintain a consistent translational capacity. For multi-protein complexes, the burst profiles of constituent genes must be coordinated to support their efficient assemblies and functions.

### The differential transcription burst strategies in scaffolded multi-protein complexes

We expanded our previous analysis of TNRC6A/B/C (the scaffold) and AGO1/2/3 (the active effector) of the miRISC complex to a diverse array of regulatory multi-protein complexes spanning mRNA decay, signal transduction, vesicle trafficking, and synaptic organization. Our analysis reveals a common transcriptional strategy that partitions proteins by their functional roles. We demonstrate that structural scaffolds are “hardwired” for reliability, utilizing a high-frequency, small-burst-size kinetics to maintain a steady, low cell-to-cell noise baseline. In contrast, their active effectors are built for responsiveness, utilizing infrequent but massive transcriptional bursts. This bifurcated strategy suggests that evolution differentially prioritizes the stable availability for the “structural chassis” (the scaffold) versus the metabolic efficiency for its cargo (the effector partners).

### Functional implication for the operation of scaffolded protein complexes

These results contribute to our understanding of the operation of scaffolded multi-protein complexes. As structural backbones that physically tether, organize, and position multiple interacting proteins, scaffolds coordinate biochemical events with spatiotemporal precision and specificity^4^. Mathematical modeling of scaffolds, such as those in the MAP kinase cascades^36^, suggests that the scaffold often acts as the rate-limiting template for assembly. High cell-to-cell variance in scaffold expression would result in frequent detrimental stoichiometric imbalances: (1) over-saturating scaffold expression levels in a sub-population of cells can lead to effector inhibition (the “prozone effect” or squelching)^37^ and the accumulation of “empty” scaffolds; or (2) in another sub-population, scaffold expression levels can be so low that free-floating unregulated enzymes (like GSK3B or MAP kinases) are left without a docking station. Such imbalances would inherently degrade signaling fidelity and increase the risk of off-target activity. Our findings suggest that high-frequency transcriptional bursting serves as a primary noise-reduction mechanism to ensure stoichiometric reliability, maintaining the scaffold at a concentration that effectively “buffers” the more volatile fluctuations of its enzymatic partners.

*Summary*. Our study establishes a fundamental link between a protein’s physical role within a multi-protein regulatory complex and the transcriptional burst kinetics of its gene. Across diverse biological systems—from the synaptic density to the WNT signaling pathway— structural scaffolds consistently utilize high-frequency, small-sized bursts to suppress cell-to-cell noise. This “scaffold law” ensures the stable availability of architectural templates, preventing the signaling failures associated with stoichiometric imbalance. These findings suggest that promoter architectures have been evolutionary tuned not just for expression abundance, but for the specific spatiotemporal demands of the macromolecular machines they constitute.

## Acknowledgement

We acknowledge Dr. Muchun Niu of Baylor College of Medicine for critical suggestions.

